# Novel Magnetic Resonance Imaging strategy targeting Neurotensin Receptors in detection of Prostate Cancer

**DOI:** 10.1101/180042

**Authors:** Mitul Desai, Neville N.C. Tam, Robert. Carraway, Shuk-Mei Ho, Craig. Ferris, Jean. King

## Abstract

Prostate cancer is the second leading cause of all male cancer deaths. One of the factors present in malignant prostate cells and shown to support its metastatic growth is the neuropeptide neurotensin (NT). The primary goal of the present study was to establish the feasibility of using a newly developed paramagnetic receptor ligand for NT and non-invasive ultrahigh-field magnetic resonance (MR) imaging to visualize prostate cancer in rodents. Orthotropic xenografts were initiated in six-week old male BALB/c *nu/nu* athymic mice (n = 28) by intra-prostatic (ventral lobe) inoculation of human prostate cancer cells (10μL of PC3 cells (10^6^ /100μl)). Palpable tumors developed within 30-60 days. A micro-imager utilized in these studies was an actively shielded 9.4T, 89 mm bore, Oxford superconducting magnet with a 100 gauss/cm gradient system. Prior to contrast injection, T2 weighted anatomy scans were done to localize the tumor with a spin-echo multi-slice sequence with TR: 2000 TE: 40 and NEX: 1 in both coronal and axial planes. The paramagnetic ligand data sets were collected with a spin-echo, T1 weighted pulse sequence (MSME): TR 300 msec; TE 5 msec; NEX 4 in both axial and coronal planes. The data sets were taken initially at 5-min intervals post contrast injection for the first half hour and then at 15 min intervals for the next 1.5-2 hours for a time series analyses. The temporal distribution of MR signal intensity in various regions were determined in the absence and presence of NT. Our results confirm that the novel NT molecule was protected from enzymatic degradation and capable of forming a high affinity paramagnetic NT ligand with an extended half-life. During the imaging studies, the signal intensity increased by 200 % in the region of the tumor. This increase in signal intensity approached maximum binding within 30 minutes and remained visible for 1 hour post-injection of the contrast agent. Taken together, these findings suggest that it is feasible to detect and image prostate cancer using a paramagnetic NT ligand and the emergence of the NT receptor ligand that may be used as a diagnostic marker for prostate cancer in humans.

## INTRODUCTION

Prostate Cancer is the second leading cause of death from cancer and is the most common malignancy in north-American men, with 1 in every 5 patients developing invasive cancer. It accounts for an estimated 29% of all new cancer cases diagnosed in U.S. men (1). Although the current diagnostic processes for the prostate cancer (including the digital rectal examination, measurement of prostate-specific antigen, transrectal ultrasonography and biopsy) may have increased (2) the number of men being screened, the mortality rate for this disease has not subsided. To minimize mortality, emerging methods of localized tumor therapy (aimed at replacing surgery and radiation) are being considered particularly establishing the requirement for accurate, concise 3D localization of prostate tumors.

Neurotensin (NT) is a 13 amino-acid peptide found in the central and peripheral nervous systems and endocrine cells of several tissues (3, 4). Recent evidence shows NT can promote the development and growth of certain tumors, particularly pancreatic (5, 6), lung (7), colon (8) and prostate cancers (9, 10). Blocking the NT receptor with the highly selective antagonist, SR48698 inhibits the tropic effects of NT on xenografted human colon (8, 11) and prostate cancer (9) cells (PC3). The presence of NT receptors in tumor correlates with the emergences of cancerous growth. These findings suggest that the emergence of NT receptor may be a diagnostic marker for prostate cancer. Considering the strong association of NT with prostate cancer, we have developed a paramagnetic ligand that recognizes the NT receptor.

With the advent of animal MR spectrometers of ultrahigh-field strengths like 9.4 Tesla (T) that enables to study biological structures of 50μm or less and technical developments in MRI combined with the use of selective contrast agent can potentially improve early detection, diagnosis and 3D localization of prostate tumor (12, 13, 14). This manuscript explores the viability of using the newly developed ligand (as a contrast agent) and MR Imaging to demonstrate the presence and extent of prostate cancer. The key to visualizing NT binding sites is the internalization of the receptor-ligand complex and the concentration of the paramagnetic atoms inside the cells. The presence of paramagnetic NT receptor ligand shortens T1 relaxation, thus creating a contrast in the MR Image (due to difference in T1 relaxation rates), from surrounding tissues.

MRI using gadolinium (GD) enhancement is becoming a common technique in the study of prostate cancer (15, 16). When gadolinium is combined with NT (Gd-NT), our results show that the paramagnetic ligand complex binds to the NT receptors in the tumor xenografted into the prostate of nude mice. MR Images showed enhanced signal intensity in the areas of the tumor as compared to the surrounding tissue. The use of antagonist (SR48692) illustrates that the receptor-ligand interaction exhibited saturablity (the antagonist blocks NT receptors in a dose dependent manner) (8). Our experiments also demonstrate that the newly developed Gd-NT complex shows tumor specificity, sensitivity with an enhanced half-life.

## MATERIALS AND METHODS

### Animals

These studies used six-week old male balb/c nu/nu athymic mice (nude mice) obtained from Charles Ricer Breeding Laboratory (Wilmington, MA). The mice were housed under aseptic conditions (enclosed overheated laminar hood, sterilized cages, bedding and water) and maintained at ambient temperature (22-24° C) in a light controlled (14 h/day) room. All animals were maintained in an accredited facility at University of Massachusetts Medical School. Food and water was provided *ad libitum*. All animals were acquired and cared for in accordance with the guidelines published in NIH Guide for Care and Use of Laboratory Animals (National Institutes of Health Publications N. 80-23, Revised 1996).

### Surgical Orthotropic Implantation

PC3 cells are known to be one of the most aggressive prostate cancer cell lines, which is easily xnografted into nude mice (17). Right before implantation, PC-3 cells (obtained form ATCC) were trypsinized and re-suspended in DMEM with 10% FCS. Xenografts were initiated in the mice (n=28) by intra-prostatic injection. Brief surgical procedures for transplantation of tumors were performed under aseptic conditions and the animals were anesthetized with isoflurane (1.5%). 20 μL of PC3 cells (2x10^6^/ 200 μL) were then injected into the ventral lobe of prostate, using Hamilton syringe with a 30 gauge needle (18, 19). The abdominal incisions were then sutured (with discontinuous, absorbable suture) using 6.0 nylon thread and the animals were then transferred to a recovery cage. Post surgical observations including the assessment of animals for signs of infection were carried out daily for next 14 days.

### MRI Study

The micro imager utilized in these studies is an actively shielded 9.4 T, 89 mm bore, Oxford superconducting magnet with a 100 gauss/cm gradient system (Resonance Research Inc.). Mice were implanted with tail vein for delivery of contrast agent prior to imaging study. Prior to contrast injection, T2 weighted anatomy scans were obtained to localize the tumor, with a spin-eco multi-slice sequence with TR: 2000 ms TE: 40 ms and NEX: 1 in both coronal and axial planes.

Once the tumor is localized on the MR Image the paramagnetic ligand (Gd-NT) was then injected in the mice (through the tail vein 1 ml/s). Post Gd-NT (0.1 mmol) administration data sets were collected with a multi-slice (30 slices per plane) spin-echo, T1 weighted pulse sequence: TR 300 ms; TE 5 ms; NEX 4 in both axial and coronal planes. The data sets initially were taken at a 5-min interval, for first 30-min, immediately post contrast injection and then-after at a 15-min interval for the next 1.5-2 hours for a time series analyses. The temporal distribution of MR signal intensity in various regions was determined in absence and presence of Gd-NT. Specific binding activity expressed as percent change in MR signal intensity/unit area over time. Specific binding were estimated as the difference in MR signal intensity for a given region of the tumor in the absence and presence of NT receptor ligand. Respiratory motion during scan was minimized with an abdominal belt. Care was taken during data analysis to correct for any abdominal/peristaltic movement in the 2-3 hours of scanning. In our anatomical scans tumors areas were identified as the areas with increase signal intensity on T2-weighted images and change in signal intensity, post Gd-NT administration, was followed on the T1-weighted images.

Experiments using SR48968 (antagonist) were carried out to test the saturablity of the Gd-NT ligand. Four different doses (5, 4, 3 and 1mmol) of SR48968 were administered intravenously, just prior to Gd-NT administration, but the later part of the experimental procedures was kept exactly the same as the one used for the Gd-NT ligand.

### Autopsy and histology

At the end of experiments (4-6 weeks after the tumor cells inoculation), mice were sacrificed by CO_2_ asphyxiation. Careful gross examination of the primary and secondary growth of prostate tumor was performed at autopsy. The primary tumor of the prostate, various prostatic lobes, seminal vesicles, urinary bladder and the urethra were excised from the mice. Regional and distant lymph nodes, lungs, liver, spleen, lower abdominal wall (organs where metastasis could occur) were subjected to the evaluation of metastasis. All tissues collected were fixed in 10% neutral buffered formal saline, routinely embedded, sectioned and stained with hematoxylin and eosin for standard microscopic examination.

## RESULT

### Establishment of orthotopic xenografts of human prostate cancer cells

From our previous studies on NT PC3 cells growth in vivo ectopic tumors were visible to eye by Day 15 post xenograft and significant difference in NT binding were recognized by Day 16 post xenograft. However, when the PC3 cells were implanted into the ventral prostate (orthotropic) the tumor growth was less aggressive and significantly different form that observed with ectopic tumor. In order to see the earliest formation of tumor, imaging began 72 hrs after the PC3 cells were xenografted into the nude mouse. The pilot studies demonstrated that inoculation with PC3 cells were not clearly visible with MRI until the end of the first week and were palpable 30-60 days post inoculation. Several regions of interests (ROIs) including the tumor –PC3 implanted region, bladder and the healthy prostate gland were identified and followed.

The growth of inoculated human PC3 cells in murine ventral prostate gland resulted in gross and palpable tumors accompanied by local invasion into the adjacent lateral and dorsal lobes of prostate. The regular glandular architecture of the involved prostate lobes was dislodged by the overwhelming growth of human prostate cancer cells (Fig 2[2]). In some (three) mice, tumor-infiltrated lymph nodes and lower abdominal wall were observed adjacent to the primary tumor. During the course of our study (90 days) no distant metastases were detected in visceral organs (liver, kidney, lung and spleen) and distant lymph nodes by gross and microscopic examinations. Fig 2[3] demonstrated the histological details of the poorly differentiated phenotypes of PC3 xenografts in the murine prostate gland.

### MRI and Contrast Agent

A mouse, 45 days post inoculation, was used to acquire the MR images shown in Fig. 1 A, B & C. Once the control MR images (T1 and T2 weighted images in coronal and axial plane) were acquired, 0.1mmol Gd-NT contrast agent was administered intravenously and post-contrast T1 weighted (coronal and axial) images were acquired every 5-min for first 30-min and at an interval of 15-min there after for 90-min. As shown in Fig. 1 D signal intensity enhances in the inoculated lobe of the prostate as compared to non-inoculated lobe suggesting that the ligand is site specific. Maximum binding is acquired in about 15-30 minutes and the contrast lasts for about 90 minutes post injection. In contrast the regular tissue (without tumor) dose not shows any signal intensity enhancement (Fig. 1 D).

**Figure. 1.**
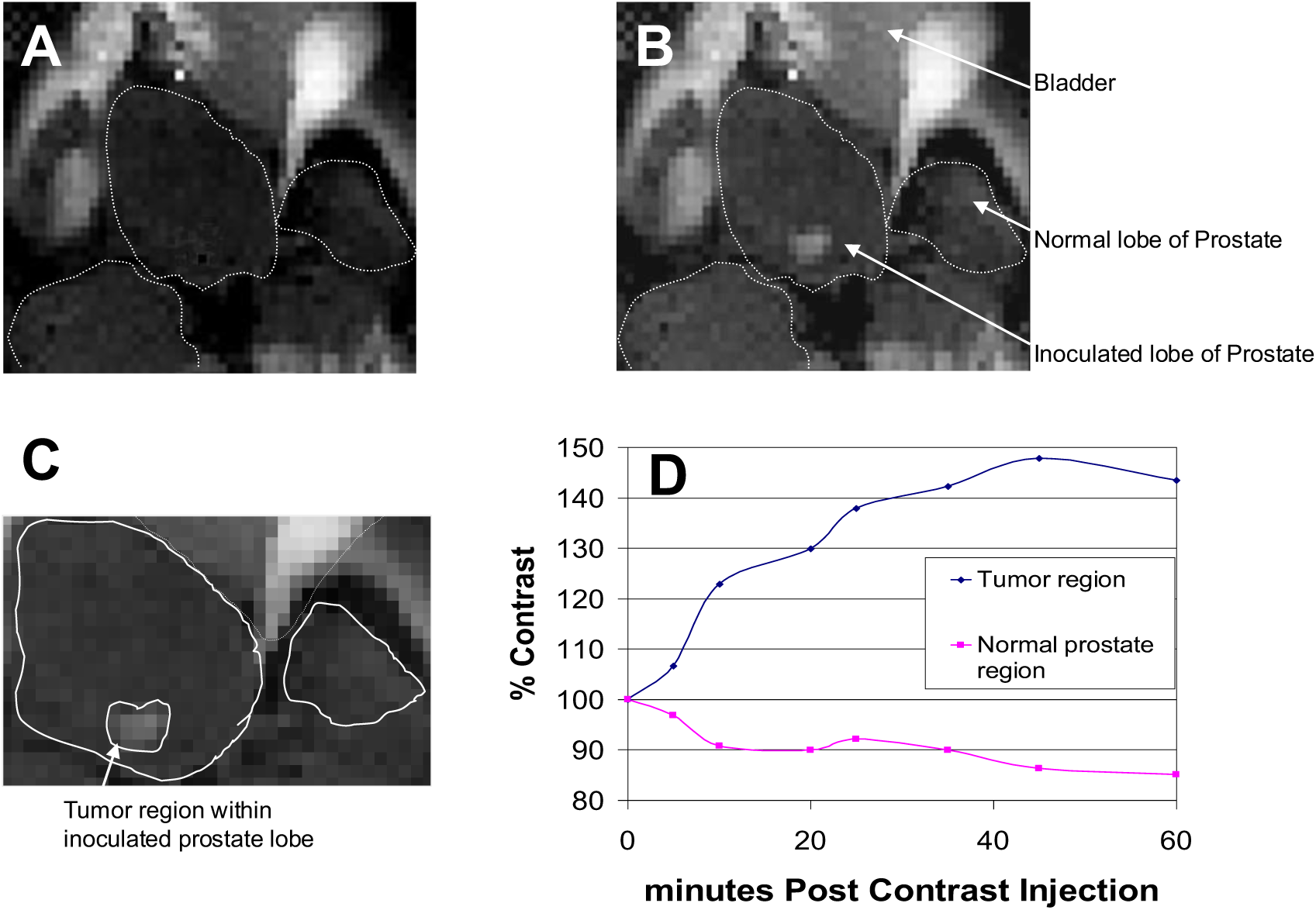
T1 (TR=300ms, TE=5ms, NEX=4) weighted MRI of prostate region showing a developed tumor and the contrast with the tumor (A and B; pre-and 15 minute post-contrast MR images. Prostate gland outlined.). A % contrast change (% of control) timeline (D) shows that the signal intensity change in tumor region is much higher as compared to control region (normal prostate).

### MRI and Histo-Patalogy

The MRI data acquired were corroborated with histological analysis. The histopathologic images confirmed the size and localization of the tumor as obtained from the MRI studies. Three axial T2 weighted (TR=2000ms, TE=40ms, NEX=1) MR Images were aligned with its corresponding histopathologic images (Fig.2 (1) & (2)). The tumor in this case grew up to 1cm in diameter (6 weeks post inoculation), which was confirmed by both MRI and histolopathologic results. It is very easy to see the tubules and the tumor cells in the high resolution histopathologic image (Fig. 2 (3) A).

**Figure. 2.**
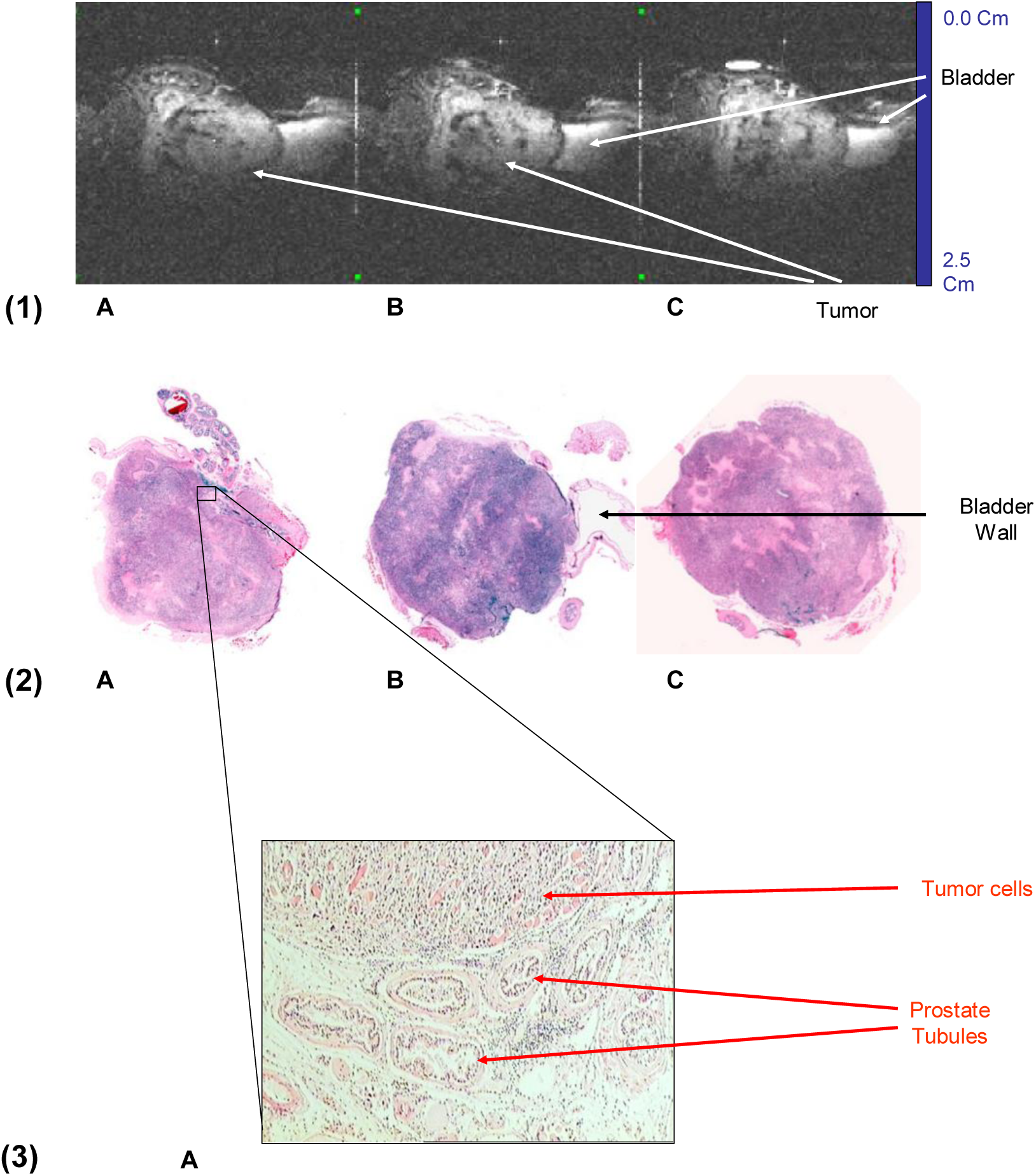
Three T2 weighted MR Images (1: A, B & C) of prostate tumor region are aligned with corresponding histopathologic slides (2: A, B & C). A high resolution histopathologic image shows tumor cells and prostate tubules.

### Sight Specificity

The MR Images collected Pre (Fig. 3 C) and 5 minutes Post (Fig. 3 D) Gd-NT administration shows that the contrast enhancement is site specific, tumor regions, and can be easily visualized. These images are spatially corroborated with histopathologic (Fig. 3 A) and T2 weighted MR image (Fig. 3 B) of the tumor region. We also found that the contrast started to develop in the bladder; this was very consistent in all of our studies. This suggests that paramagnetic ligand is site specific and has superior half life, as the contrast form all the other organ (tissues) is washed out very quickly (within 5 min) except the tumor region where the ligand is bound with the NT receptor and stays in there for as long as 90-min.

**Figure. 3.**
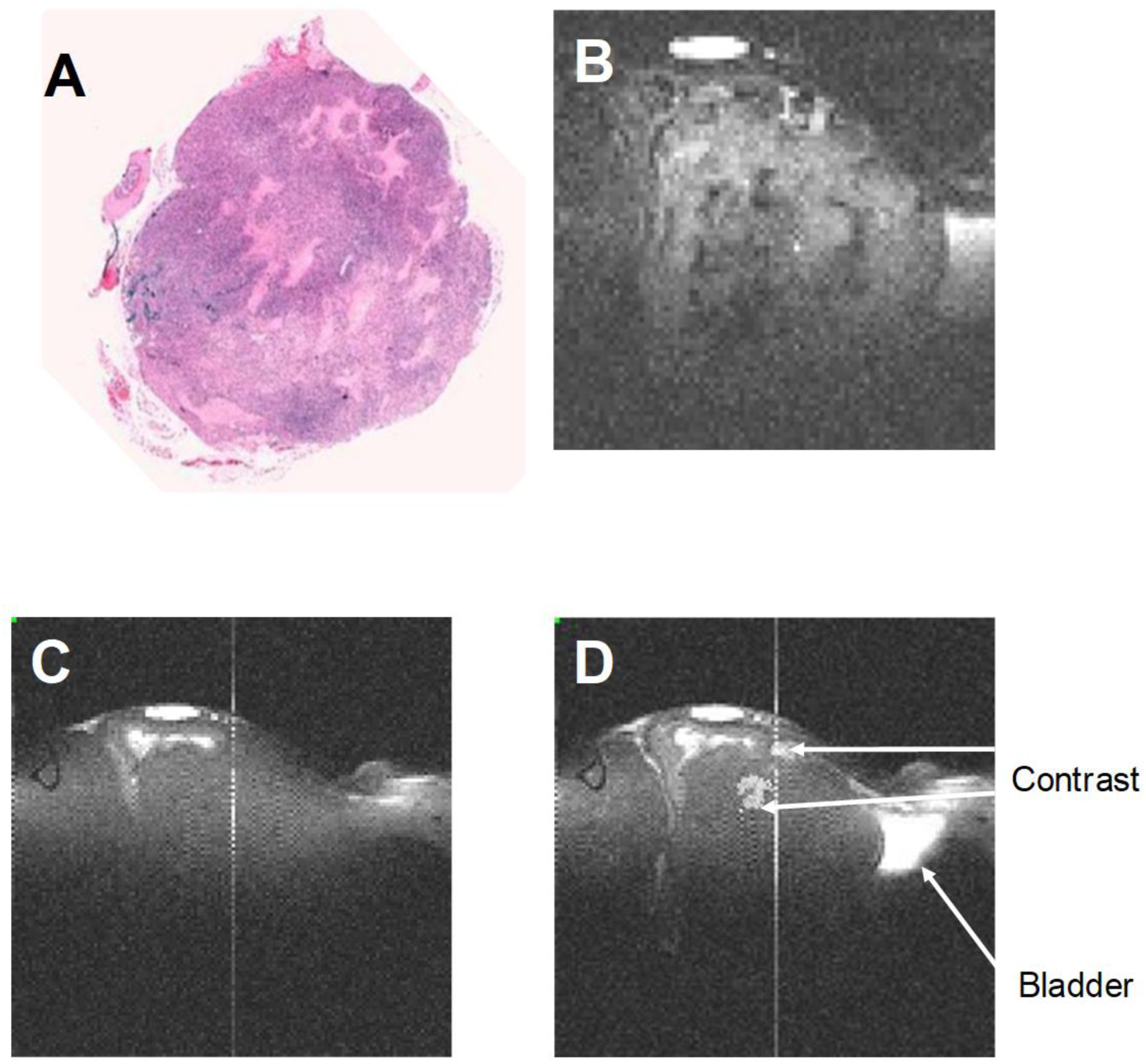
Histopathologic image (A) of prostate tumor is aligned with its corresponding T2 weighted MR Image (B). Corresponding pre (C) and 5 minutes post-contrast (D) T1 weighted images are also aligned. Post-contrast T1 weighted image shows contrast within the prostate tumor.

### Longitivity & Saturablity

A comparative study between receptor-ligand complex and antagonist (SR48698) shows that the NT receptor ligand exhibits saturablity, as seen in Fig. 4. The plot of % contrast change against time shows that there is no signal intensity enhancement in the antagonist case, where as the signal intensity enhances up to 200% and is sustained up to an hour.

**Figure. 4.**
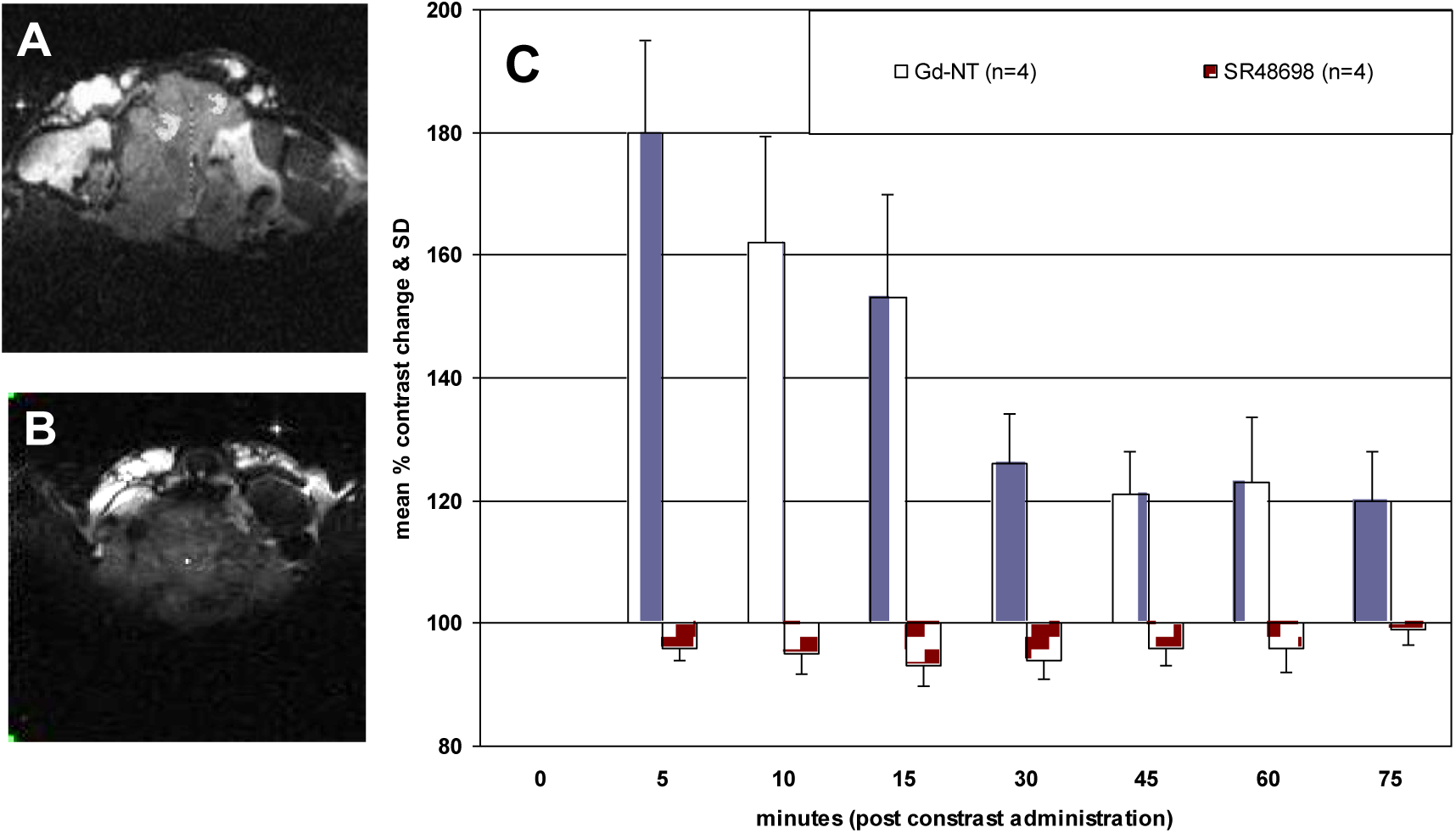
Axial T1 weighted (TR=300ms, TE=5ms, NEX=4), 10 minutes post-contrast, MR images are acquired for two cases; Gd-NT contrast agent and antagonist (A and B). Mean % contrast change and standard deviation (SD) is compared (C) for both the cases and it can be seen that the contrast stays in the tumor for about 60 min in the case of Gd-NT contrast administration. Error bars indicate SD from the mean.

## DISCUSSION/CONCLUSION

Our Studies indicate that the key to visualizing NT binding sites is the internalization of the receptor-ligand complex and the concentration of the paramagnetic atoms inside the cells. The presence of paramagnetic NT receptor ligand shortens T1 relaxation, thus creating a contrast in the MR Image (due to difference in T1 relaxation rates), from surrounding tissues.

However, couple of issues arose when we initiated the inoculation procedure including the viability of the cells in vivo, the volume of inoculants, and the placement of the cells. Initially we tried to inoculate PC3 cells S.C into the right flank of mice midway between the arm and the leg. After many attempts, we arrived at our final protocol which decreased the volume to 20μL (lower then we initially attempted) and introduced the cells directly into the ventral lobe of the prostate. It appears that limiting the volume of cells decreased cell loss to the surrounding tissue. Although the isotropic model was appreciably more difficult to attain, it was undoubtedly more akin to the human model (18, 20, 21). These modifications currently produce much higher rate of successful tumor growth (65-70 %). The tumor grew anywhere form 1.0-1.5 cm in size after about 30-60 days post inoculation.

Although these large tumors where easily visible, it was not easy to trace small tumors using the T2 weighted images (22, 23). Use of selective contrast agent proved to be a very precise method in tracking such small tumor. The SNR (signal to noise ratio) also greatly improved with the use of contrast agent. The specificity of the paramagnetic ligand, unlike T2 weighted technique, increases the signal intensity in the tumor region and the tumor region only. At its peak the signal intensity increased to 200% of the signal intensity before Gd-NT administration, and stayed within the tumor region for as long as 30-min, making the estimation and detection process convenient and highly precise.

Pre-saturation of NT receptor with 4 different dose of antagonist SR48692 changed the MR signal intensity and there by demonstrating the dose dependent nature of the receptor-ligand saturablity.

MRI using gadolinium (GD) enhancement is becoming a common technique in the study of prostate cancer (15, 16). When gadolinium is combined with NT (Gd-NT), our results show that the paramagnetic ligand complex binds to the NT receptors in the tumor xenografted into the prostate of nude mice. MR Images showed enhanced signal intensity in the areas of the tumor as compared to the surrounding tissue. The use of antagonist (SR48692) illustrates that the receptor-ligand interaction exhibited saturablity (the antagonist blocks NT receptors in a dose dependent manner) (8). Our experiments also demonstrate that the newly developed Gd-NT complex shows tumor specificity, sensitivity with an enhanced half-life.

In summary, we have shown that the MR images exhibited high contrast enhancement of tumor region and tumor region only, post Gd-NT ligand administration. This study supports the use of Gd-NT ligand in conjunction with non-invasive Magnetic Resonance Imaging as a useful tool for early detection, diagnosis and 3D localization of prostate cancer.

## REFERENCES

1. Greenlee RT, Murray T, Bolden S, Wingo PA. Cancer statistics, 2000. CA Cancer J Clin 2000;50(1):7–33.

2. Kurhanewicz J, Swanson MG, Nelson SJ, Vigneron DB. Combined magnetic resonance imaging and spectroscopic imaging approach to molecular imaging of prostate cancer. J Magn Reson Imaging 2002;16(4):451–63.

3. Brawley OW, Thompson IM. Chemoprevention of prostate cancer. Urology 1994;43(5):594–9.

4. Reinecke M. Neurotensin. Immunohistochemical localization in central and peripheral nervous system and in endocrine cells and its functional role as neurotransmitter and endocrine hormone. Prog Histochem Cytochem 1985;16(1):1–172.

5. Ishizuka J, Townsend CM, Jr., Thompson JC. Neurotensin regulates growth of human pancreatic cancer. Ann Surg 1993;217(5):439–45.

6. Iwase K, Evers BM, Hellmich MR, Kim HJ, Higashide S, Gully D, et al. Inhibition of neurotensin-induced pancreatic carcinoma growth by a nonpeptide neurotensin receptor antagonist, SR48692. Cancer 1997;79(9):1787–93.

7. Davis TP, Burgess HS, Crowell S, Moody TW, Culling-Berglund A, Liu RH. Beta-endorphin and neurotensin stimulate in vitro clonal growth of human SCLC cells. Eur J Pharmacol 1989;161(2-3):283–5.

8. Maoret JJ, Anini Y, Rouyer-Fessard C, Gully D, Laburthe M. Neurotensin and a non-peptide neurotensin receptor antagonist control human colon cancer cell growth in cell culture and in cells xenografted into nude mice. Int J Cancer 1999;80(3):448–54.

9. Seethalakshmi L, Mitra SP, Dobner PR, Menon M, Carraway RE. Neurotensin receptor expression in prostate cancer cell line and growth effect of NT at physiological concentrations. Prostate 1997;31(3):183–92.

10. Sehgal I, Powers S, Huntley B, Powis G, Pittelkow M, Maihle NJ. Neurotensin is an autocrine trophic factor stimulated by androgen withdrawal in human prostate cancer. Proc Natl Acad Sci U S A 1994;91(11):4673–7.

11. Gully D, Canton M, Boigegrain R, Jeanjean F, Molimard JC, Poncelet M, et al. Biochemical and pharmacological profile of a potent and selective nonpeptide antagonist of the neurotensin receptor. Proc Natl Acad Sci U S A 1993;90(1):65–9.

12. Coakley FV, Qayyum A, Kurhanewicz J. Magnetic resonance imaging and spectroscopic imaging of prostate cancer. J Urol 2003;170(6 Pt 2):S69–75; discussion S75-6.

13. Claus FG, Hricak H, Hattery RR. Pretreatment evaluation of prostate cancer: role of MR imaging and 1H MR spectroscopy. Radiographics 2004;24 Suppl 1:S167–80.

14. Jacobs MA, Windham JP, Soltanian-Zadeh H, Peck DJ, Knight RA. Registration and warping of magnetic resonance images to histological sections. Med Phys 1999;26(8):1568–78.

15. Gossmann A, Okuhata Y, Shames DM, Helbich TH, Roberts TP, Wendland MF, et al. Prostate cancer tumor grade differentiation with dynamic contrast-enhanced MR imaging in the rat: comparison of macromolecular and small-molecular contrast media--preliminary experience. Radiology 1999;213(1):265–72.

16. Larson BT, Collins JM, Huidobro C, Corica A, Vallejo S, Bostwick DG. Gadolinium-enhanced MRI in the evaluation of minimally invasive treatments of the prostate: correlation with histopathologic findings. Urology 2003;62(5):900–4.

17. Lim DJ, Liu XL, Sutkowski DM, Braun EJ, Lee C, Kozlowski JM. Growth of an androgen-sensitive human prostate cancer cell line, LNCaP, in nude mice. Prostate 1993;22(2):109–18.

18. Rembrink K, Romijn JC, van der Kwast TH, Rubben H, Schroder FH. Orthotopic implantation of human prostate cancer cell lines: a clinically relevant animal model for metastatic prostate cancer. Prostate 1997;31(3):168–74.

19. Wang X, An Z, Geller J, Hoffman RM. High-malignancy orthotopic nude mouse model of human prostate cancer LNCaP. Prostate1999;39(3):182–6.

20. Thalmann GN, Anezinis PE, Chang SM, Zhau HE, Kim EE, Hopwood VL, et al. Androgen-independent cancer progression and bone metastasis in the LNCaP model of human prostate cancer. Cancer Res 1994;54(10):2577–81.

21. Thalmann GN, Sikes RA, Wu TT, Degeorges A, Chang SM, Ozen M, et al. LNCaP progression model of human prostate cancer: androgen-independence and osseous metastasis. Prostate 2000;44(2):91–103 Jul 1;44(2).

22. Schiebler ML, Tomaszewski JE, Bezzi M, Pollack HM, Kressel HY, Cohen EK, et al. Prostatic carcinoma and benign prostatic hyperplasia: correlation of high-resolution MR and histopathologic findings. Radiology 1989;172(1):131–7.

23. Berman C, Brodsky NJ. Prostate Cancer Imaging. Cancer Control 1998;5(6):541–54.

